# Protein co-scientist aided simulations of the structural and functional impact of O-GlcNAc Transferase TPR domain deletions and amino acid substitutions associated with X-Linked Intellectual Disability

**DOI:** 10.64898/2026.10.29.762748

**Authors:** Sonja Barth, Allison Bechard, Amelia Krafft, Chakkai Yip, Caleb Aboagye, Charlie Fehl, Geoffrey H. Siwo

## Abstract

O-GlcNAc transferase (OGT) is an essential glycosyltransferase that regulates cellular processes through the addition of O-linked β-N-acetyglucosamine (O-GlcNAc) to target proteins. Substrate recognition and protein-protein interactions are mediated by the tetratricopeptide repeat (TPR) domain, yet the contributions of individual TPR repeats to structural stability and function remain poorly understood. Here, we employed an integrated computational framework combining an AI protein co-scientist, Speak to a Protein, which incorporates Boltz-2 for structure prediction and rDock for ligand binding evaluation, with external tools including AlphaFold3 for more advanced structure prediction and the HDOCK server for protein-protein docking. This framework was used to investigate how deletions of TPR domains and disease-associated amino acid substitutions within the TPR domain affect OGT structure and function. Five synthetic truncation OGT isoforms, thirteen single-TPR deletion variants, and five X-linked intellectual disability (XLID)-associated amino acid substitutions were modeled and aligned to wild-type OGT for comparison. Structural perturbations were quantified using backbone root mean square deviation (RMSD), while functional effects were assessed through docking analyses with UDP-GlcNAc, Host Cell Factor 1 (HCF1), and Tet methylcytosine dioxygenase 1 (TET1). Variation in TPR length altered global protein structure and induced long-range conformational changes within the catalytic domain despite no sequence modifications in that region. Single TPR deletions showed that individual repeats contribute unequally to structural stability, with deletions in mid-to-late TPR domains producing the largest structural disruptions. Disease-associated amino acid substitutions also generated conformational changes and reduced predicted ligand binding, consistent with known reductions in enzymatic activity in XLID. Protein-protein docking analyses demonstrated substrate-specific effects, as domain deletion and amino acid substitution variants of OGT had different interactions with TET1 and HCF1. Collectively, these findings identify the TPR domain as a regulatory domain that governs structure, ligand recognition, and protein interactions. More broadly, this work demonstrates how the integration of a protein co-scientist with protein structure modeling tools can be used to predict the structural and functional consequences of domain deletions or natural amino acid variants.

## Introduction

O-GlcNAc transferase (OGT) is an essential mammalian enzyme that catalyzes the addition of O-linked β-N-acetyglucosamine (O-GlcNAc) to serine and threonine residues^1^. OGT is required for cell proliferation in mammals, glycosyltransferase activity, and cleavage of Host Cell Factor 1 (HCF1), an important transcriptional coregulator involved in cell-cycle regulation^2^. Because OGT modifies many nuclear and cytoplasmic proteins, disruption of its activity can affect a wide range of cellular processes. Consequently, dysregulation of OGT has been implicated in a variety of diseases, including cancer, neurodegeneration, diabetes, and X-linked intellectual disability (XLID)^3^.

Structurally, OGT contains an N-terminal tetratricopeptide repeat (TPR) domain and a C-terminal catalytic domain. While the catalytic domain is essential for enzymatic activity, the TPR domain mediates protein-protein interactions and substrate recognition. The TPR region forms a superhelical scaffold that guides substrates toward the catalytic site and binding partners for efficient modifications. Previous studies have demonstrated that alterations in TPR architecture can significantly impact OGT function^5^. Naturally occurring isoforms vary in TPR repeat number, and engineered truncation isoforms exhibit reduced glycosylation activity and impaired cellular viability, particularly when fewer than eight TPR domains are retained^4,5^. These findings suggest that individual TPR domains make distinct contributions to structural stability and substrate recognition^6^. However, the importance of specific repeats remains poorly understood.

XLID is frequently caused by mutations within the TPR domains, which lead to HCF1 misprocessing, OGT misfolding, and downstream neurological abnormalities^3^. These observations suggest that the pathogenic effects of OGT variants may arise not only from loss of catalytic activity but also from altered protein conformation and disrupted interactions with key substrates. Despite growing recognition of the TPR domain as critical for OGT function, the structural consequences of specific domain deletions and disease-associated mutations remain incompletely characterized.

Recent advances in artificial intelligence-based protein structure predictions provide an opportunity to investigate the relationship between TPR architecture and OGT function systematically. Computational approaches can complement traditional biochemical studies by enabling large-scale evaluation of structural perturbations that would be difficult to assess experimentally^7^. In the present study, we implemented deep learning-based structural modeling to characterize naturally occurring OGT isoforms, single-TPR deletion variants, and XLID-associated amino acid substitutions. Predicted structures were analyzed to quantify changes in global conformation, local stability, and catalytic domain positioning relative to wild-type OGT.

To further assess the functional consequences of structural variations, we examined interactions between OGT variants and three biologically relevant binding partners. UDP-GlcNAc, the native glycosyl donor, was selected because of its central role in OGT-mediated glycosylation. HCF1, a canonical OGT substrate, was included due to its essential role in transcriptional regulation^9^. In addition, Tet methylcytosine dioxygenase 1 (TET1), an OGT-interacting protein involved in chromatin regulation, was used to evaluate changes in protein-protein interaction potential^10^.

Together, these targets enable evaluation of how TPR domain alterations influence the catalytic domain, substrate recognition, and protein-protein interactions. Docking analyses were therefore performed using predicted OGT structures to compare the interaction stability and binding affinity across naturally occurring isoforms, engineered deletion variants, and disease-associated amino acid substitutions.

By integrating AI-driven structural prediction with docking interaction analysis, this study establishes a predictive framework for understanding how alterations in TPR architecture influence OGT structure and function. We analyzed synthetic OGT isoforms, single-TPR deletion variants, and XLID-associated missense mutations to determine how changes to the TPR domain influence global protein architecture, catalytic domain stability, ligand binding, and protein-protein interactions. We hypothesized that individual TPR repeats contributed unequally to OGT structure and that targeted deletions within specific repeats would selectively alter substrate recognition and interaction networks. By systematically comparing specific TPR deletions and disease-associated variants, this work demonstrates the potential for AI co-scientists in investigating the impact of domain deletions and amino acid variants.

## Materials and Methods

A key innovation of this work is the use of Speak to a Protein from PlayMolecule.org, a multimodal natural-language-driven AI co-scientist optimized for interactive protein structure analysis in combination with protein structure prediction tools, to predict the structures of synthetic TPR-truncated isoforms, single-TPR deletion variants, and disease-associated amino acid substitutions^8^. These predicted models enable systematic comparison of how changes in TPR length or point mutations alter overall OGT conformation and catalytic-domain positioning. By aligning each predicted structure to wild-type OGT and evaluating structural metrics, we can systematically identify which TPR regions are most critical for maintaining structural integrity and functional stability. In addition to structural analysis, we evaluate how these perturbations influence OGT interactions with key substrates.

### Protein Sequence Design

All variant protein sequences were derived from the human wild-type OGT sequence (UniProt accession O15294). In our first experiment, to evaluate how variation in TPR domain length influences OGT structure and function, five previously generated synthetic isoforms (OGT-4 through OGT-8) were selected based on previous in vitro studies examining cell growth and HCF-1 binding, to compare with in silico results^5^. Isoform structures were generated in silico using the Boltz2^19^ modeling framework via the protein co-scientist platform - Speak to a Protein^8^. Each variant retained the full C-terminal catalytic domain while differing in the number of N-terminal domains removed (Figure 1). OGT-4 removed 9 TPR domains and was the shortest, while OGT-8 removed 5 and was the longest.

**Figure 1:**
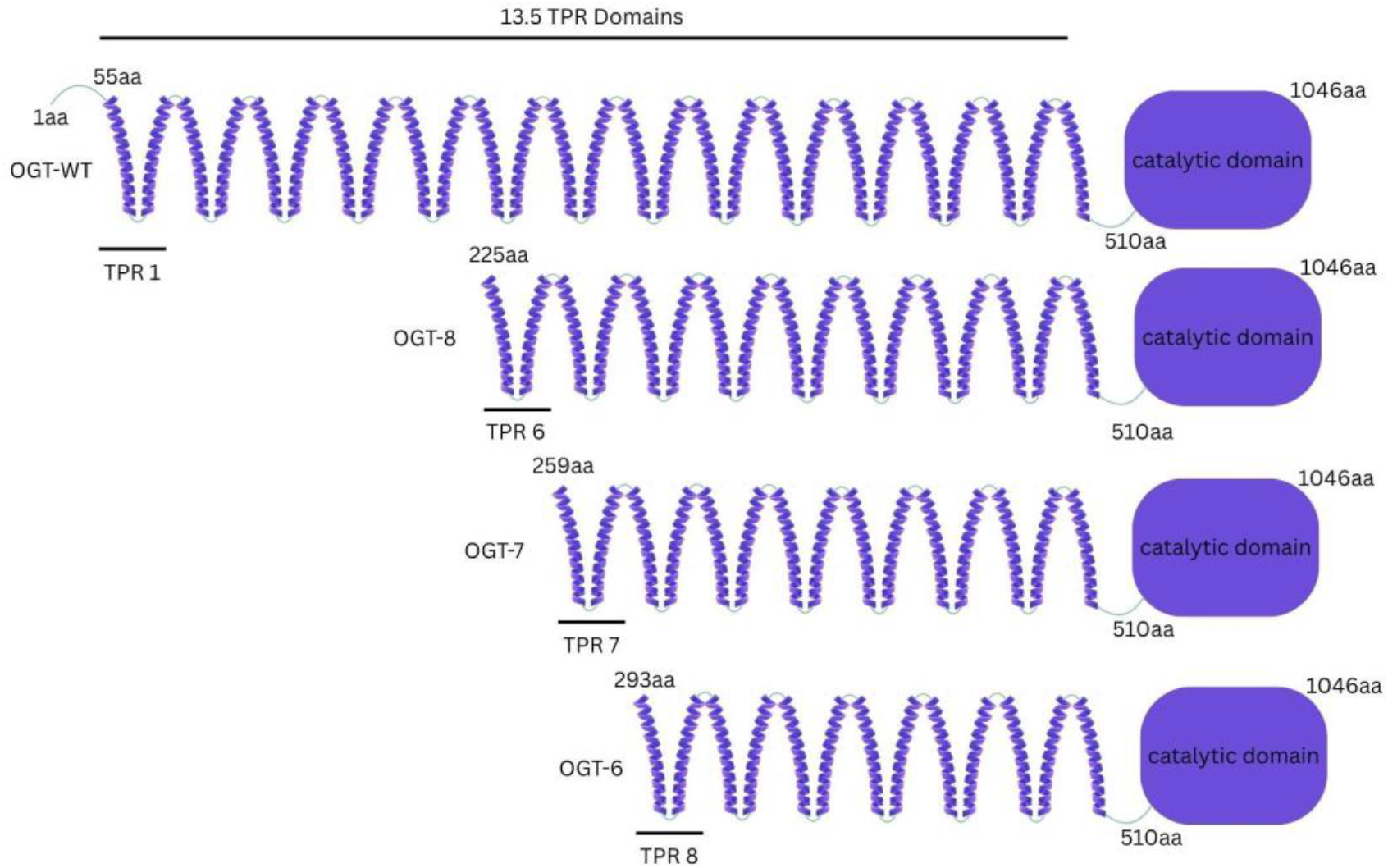
Wild-type OGT (OGT-WT) contains all 13 TPRs followed by the C-terminal catalytic domain (∼aa 510–1046). Truncated isoforms (OGT-4 through OGT-8) progressively remove TPRs 6-13, generating shorter constructs that retain the full catalytic domain but differ in substrate-binding architecture^6^.

Next, to determine how individual TPR domains contribute to OGT structural stability and function, thirteen single deletion variants (delTPR1-delTPR13) were generated in silico from the wild-type OGT sequence by removing one TPR domain at a time using Speak to a Protein (Figure 2). Each variant retained the N-terminus and full catalytic domain, with only the specified TPR domain removed. The residues flanking each deleted region were directly joined without introducing any linker residues to the protein sequence.

**Figure 2:**
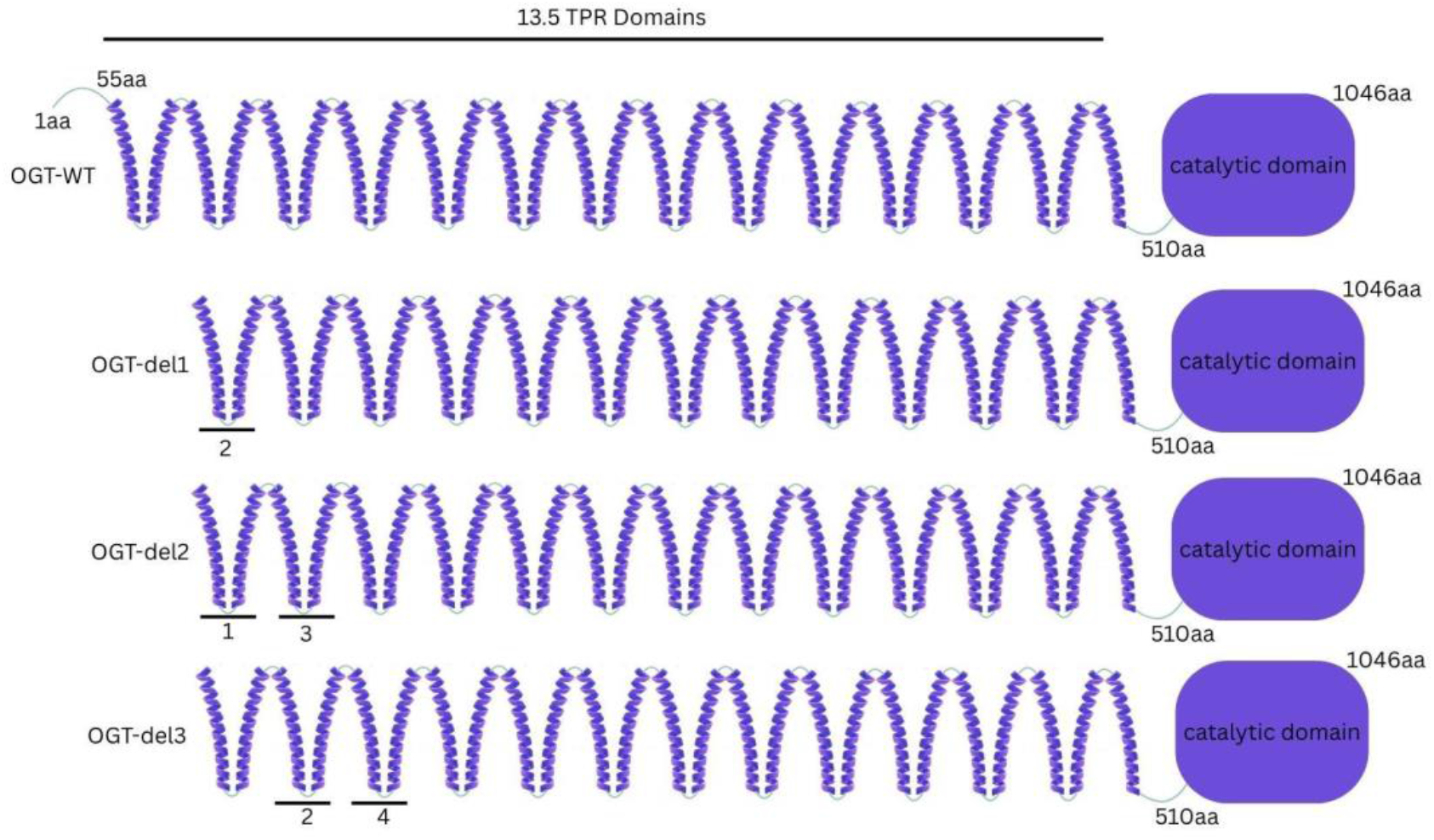
Wild-type OGT (OGT-WT) contains all 13 TPRs followed by the C-terminal catalytic domain (∼aa 510– 1046). Truncated isoforms were generated that remove each TPR domain. OGT-del1 removes TPR domain 1, OGT-del2 removes TPR domain 2, and OGT-del3 removes TPR domain 3.

Then, to assess the structural and functional consequences of disease-associated mutations, five missense mutations associated with XLID were selected based on prior literature: L254F, A259T, R284P, A319T, and E339G^3^. All mutations were located within the TPR domain and introduced by a single amino acid substitution in the wild-type sequence while keeping the rest of the sequence unchanged.

Lastly, to evaluate how structural perturbations in OGT influence protein-protein interactions, delTPR1, delTPR3, delTPR5, and OGT-8 were chosen for further analysis of binding with HCF1 and TET1. TET1 has been shown to have lower binding affinity when the TPR domain region 7-13.5 is deleted^12^. Additionally, these selected point mutation variants affect both HCF1 and TET1 binding, occurring in TPR domains 6-9^11,12^. Based on this knowledge, isoforms delTPR1, delTPR3, and delTPR5 were chosen to test, as they kept all preserved key domains and didn’t have large overlaps to cover a broader spectrum of domain deletion. Additionally, based on this and previous in vitro studies of OGT isoforms, we chose to test OGT-8 against HCF1 and TET1^9^.

### Structure Prediction

Three-dimensional structures of OGT isoforms (OGT-4, OGT-5, OGT-6, OGT-7, OGT-8) and wild-type OGT were generated in silico. Wild-type OGT, containing all 13.5 TPRs, was included as the reference structure for subsequent structural comparisons and docking analyses.

Initial structure prediction was performed using the Boltz2 modeling framework via the Speak to a Protein platform. However, persistent errors within Speak to a Protein prevented reliable generation of three-dimensional structures for all variant sequences. Therefore, to maintain consistency across the final data set, all OGT isoforms, variants, and wild-type sequences were subsequently modelled on the AlphaFold-3 Server. The resulting predicted structures were then downloaded in CIF format and re-imported into Speak to a Protein for structural alignment and analysis.

### Structural Alignment

Using Speak to a Protein, predicted structures were aligned to the wild-type OGT using backbone Cα alignment, and structural deviation was quantified using Backbone RMSD for the global structure, N-terminal TPR region (residues 55-496), and C-terminal catalytic domain (residues 510-1046). Confidence in generated protein structures was assessed using predicted Local Distance Difference Test (pLDDT) with scores >90 regarded as high confidence^20^.

### Ligand Docking

To assess whether structural changes impacted ligand binding, UDP-GlcNAc, the glycosyl donor of OGT, was docked to each variant and wild-type OGT structure using Speak to a Protein. Speak to a Protein evaluates predicted ligand binding using rDock^17^ scoring. For each complex, the top 20 docking poses were generated, with each pose representing alternative predicted binding conformations. rDock scores were obtained for each pose and used to compare predicted binding across OGT structures, with more negative scores indicating more favorable predicted interactions ^17^. Docking score distributions were summarized across the 20 poses and visualized using the box plot feature in GraphPad Prism. Wild-type OGT was used as the control for all structural and docking analyses.

### Protein-Protein Docking

To assess whether structural changes impacted large molecule binding, HCF1 and TET1, known protein interaction partners of OGT, were independently docked to each OGT variant and wild-type OGT structure. Human HCF1 and TET1 amino acid sequences were obtained from UniProt in FASTA format (HCF1, UniProt accession P51610; TET1, UniProt accession Q8NFU7). The FASTA sequences were submitted to AlphaFold-3^20^ for three-dimensional structure prediction, and the resulting structures were downloaded in CIF format. Since the docking tools available through Speak to a Protein were limited to small-molecule protein docking, protein-protein docking was performed using the HDOCK server^18^, a dedicated platform for protein-protein docking. The AlphaFold predicted structures of HCF1 and TET1 were independently docked to each OGT variant and wild-type OGT structure using HDOCK. For each complex, the top 20 docking poses were retained for analysis. These poses represent alternative binding configurations and correspond to an HDOCK score, which was extracted for each pose and used to compare predicted interaction favorability. Docking score distributions were summarized across the 20 poses and visualized using the box plot feature in GraphPad Prism. Wild-type OGT was used as the control for all structural and docking analyses.

### Statistical Analysis

Statistical analysis was performed using pairwise two-sample t-tests to compare rDock and HDOCK distributions between each variant and wild-type OGT. Statistical significance was defined as p < 0.05 (*), p < 0.01 (**), and p < 0.001 (***).

## Results

### N-terminal Truncations of OGT Alter Global Protein Structure

Both Boltz2 and AlphaFold3 generated high quality wild-type OGT structures, with pDLLT of 93.7 and 93.4, respectively. These high confidence scores supported the use of the predicted structures for following structural comparisons. To investigate the effects of TPR domain length on OGT conformation five previously reported synthetically truncated isoforms of OGT (OGT-4 through OGT-8) were modeled and aligned to wild-type OGT^5^. Global backbone root mean square deviation (RMSD) values were calculated to quantify structural differences between each isoform and wild-type OGT. OGT-4 and OGT-8 exhibited the largest deviations from wild-type with RMSD values approaching 1.7 Å, while intermediate isoforms showed smaller RMSD values ranging from approximately 0.5-0.8 Å (Figure 3a).

**Figure 3:**
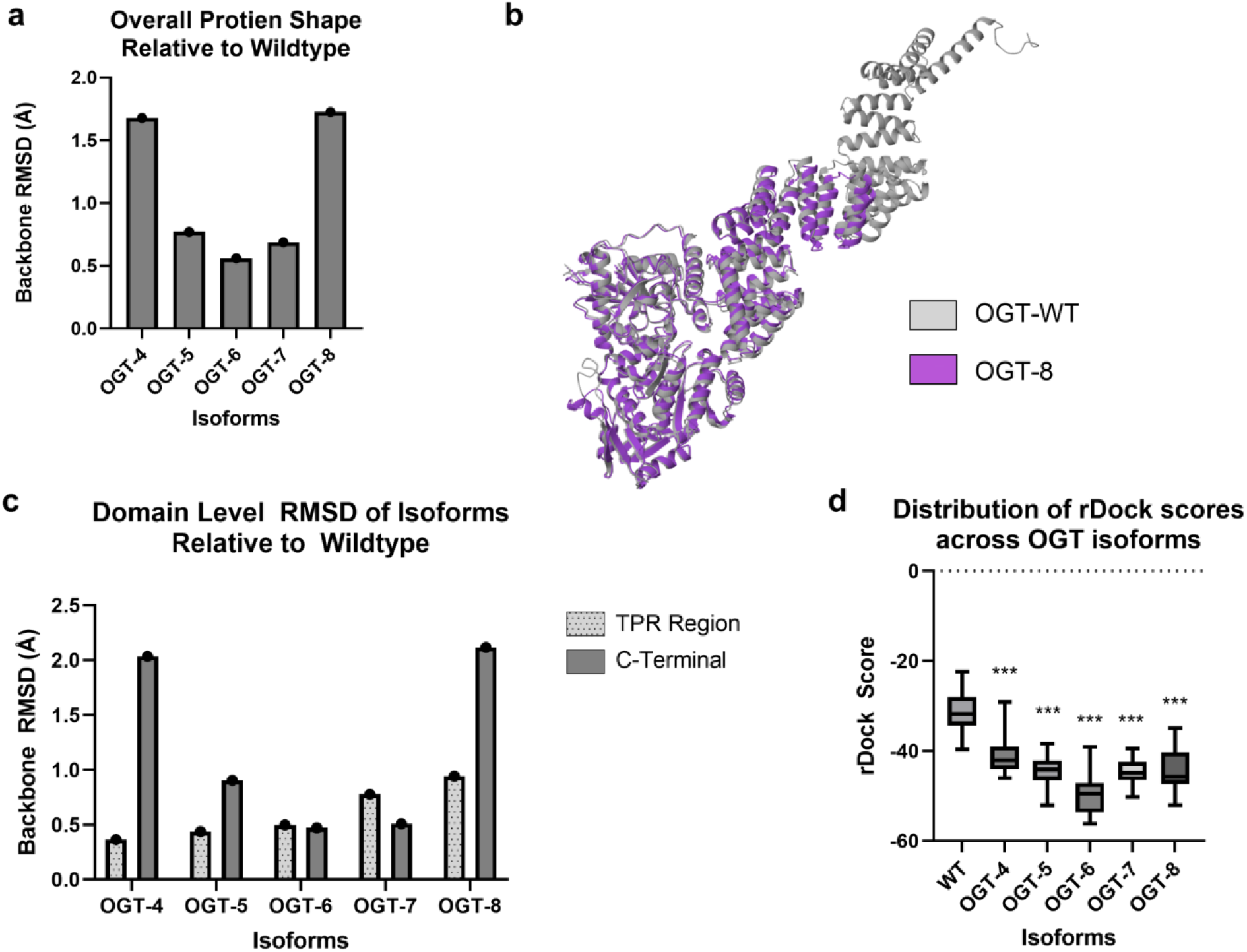
(a) Overall backbone RMSD of OGT isoforms relative to wild type. Lower RMSD indicates greater structural similarity to wild-type. (b) OGT-WT and OGT-8 aligned by the catalytic domain. (c) Backbone RMSD values for the TPR region and C-terminal catalytic domain of each isoform relative to wild-type. (d) Box plots showing rDock scores for 20 docking poses (n = 20) of UDP-GlcNAc with wild-type OGT and isoforms. More negative scores indicate stronger predicted binding. *p < 0.05, ***p < 0.001

To determine whether structural perturbations were localized within the TPR region or propagated throughout the protein, domain-specific RMSD values were calculated for the TPR superhelix and the C-terminal catalytic domain. OGT-4, OGT-5, and OGT-8 had greater structural deviations within the catalytic domain than within the TPR region, despite modifications occurring within the N-terminal TPR domain (Figure 3b). Notably, OGT-4 and OGT-8 had a catalytic domain RMSD value approaching 2.0 Å. In contrast, OGT-6 showed similar RMSD values for the TPR and catalytic regions, while OGT-7 had greater deviation within the TPR region (∼0.8 Å) than the catalytic domain (∼0.5 Å). These results demonstrate that sequential TPR deletions did not produce a uniform pattern of structural deviation across isoforms, with some truncations associated with structural differences extending into the catalytic domain and others producing more localized changes.

Structural changes were accompanied by altered ligand-binding behavior. Docking simulations revealed that all the synthetic truncation variants exhibited more favorable predicted UDP-GlcNAc binding than wild-type OGT, as indicated by more negative rDock scores (Figure 3c). Among the isoforms, OGT-6 demonstrated the strongest predicted binding affinity, whereas wild-type OGT displayed the broadest distribution of docking scores. Together, these results suggest that modification in the TPR domain length influences both OGT structural organization and ligand recognition.

### Individual TPR Repeats Contribute Unequally to Structural Stability and Ligand Binding

Deletions of individual TPR domains produced larger structural perturbations than those observed among truncated variants of OGT. Global backbone RMSD values ranged from approximately 6.5 to 11.2 Å relative to wild-type OGT, indicating that removal of a single repeat disrupts the overall structure of the TPR superhelix (Figure 4a). The magnitude of structural disruption depended on the location of the deleted repeat. Variants lacking early TPR domains (TPR1-TPR4) exhibited lower RMSD values, whereas deletions within later TPR domains (TPR9-TPR13) resulted in the largest global RMSD values.

**Figure 4:**
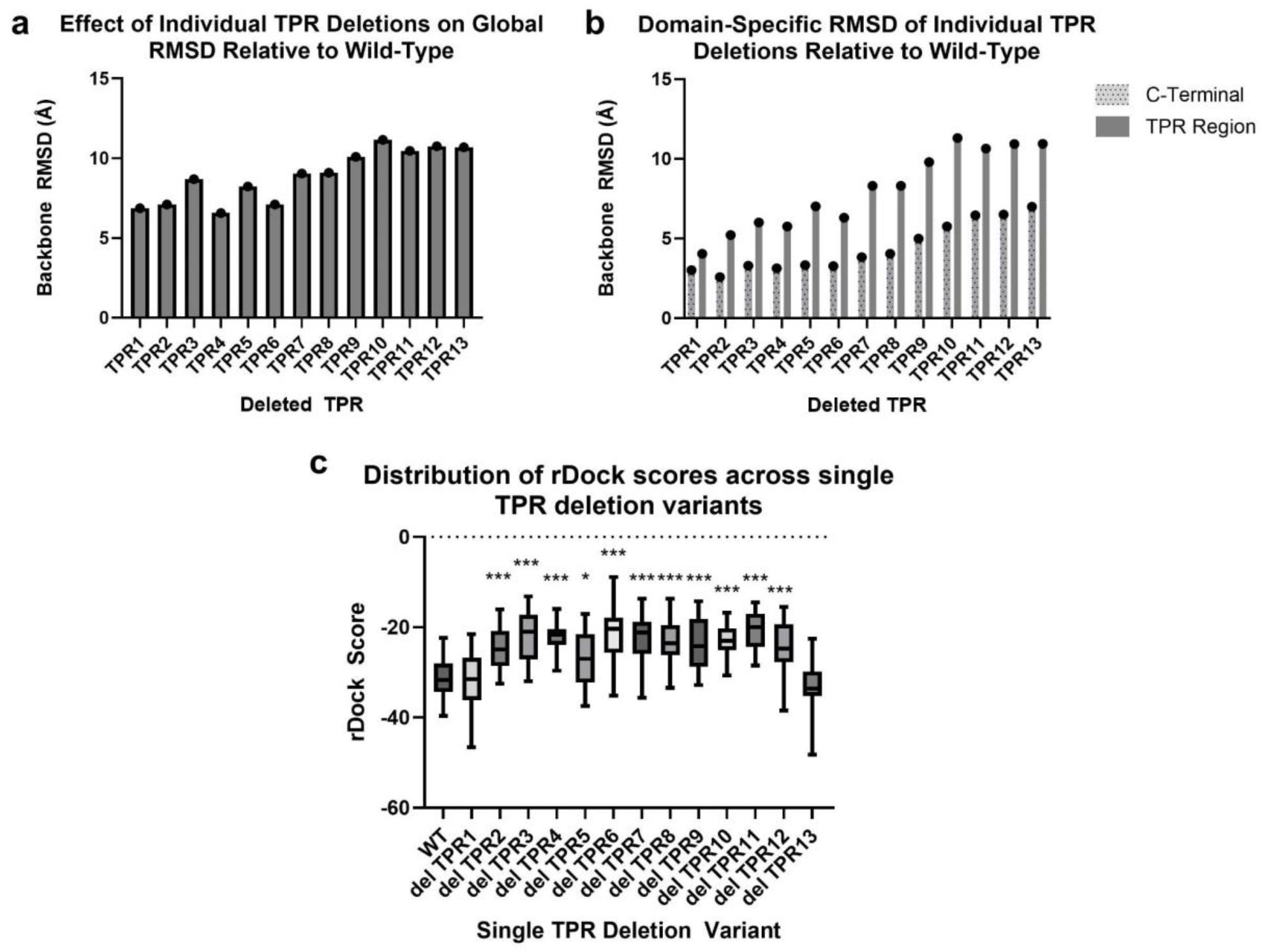
(a) Backbone RMSD values for OGT variants containing individual TPR deletions aligned to wild-type OGT. (b) Domain-specific backbone RMSD values for the TPR region and C-terminal catalytic domain of each deletion variant relative to wild-type. (c) rDock scores for 20 docking poses of UDP-GlcNAc with wild-type OGT and TPR deletion variants. More negative scores indicate stronger predicted binding. Statistical significance was determined using pairwise t-tests relative to wild-type (*p < 0.05, ***p < 0.001).

Domain-level analysis revealed that structural changes were concentrated within the TPR region but frequently propagated into the catalytic domain (Figure 4b). Across all deletion variants, RMSD values within the TPR superhelix exceeded those observed in the catalytic domain, suggesting structural changes were concentrated near the site of deletion. However, deletions within TPR9-TPR13 produced increased catalytic domain RMSD, consistent with the physical proximity of these TPR domains to the catalytic domain. These findings identify the mid-to-late TPR regions as a critical determinant of overall protein stability.

Structural disruption was accompanied by changes in predicted ligand binding. Docking simulations with UDP-GlcNAc revealed that deletions within the central TPR region (TPR4-TPR8) resulted in less negative rDock scores relative to wild-type OGT, indicating weaker predicted ligand binding (Figure 4c). In contrast, deletions near the N- and C-terminal ends of the TPR superhelix produced broader score distributions, suggesting increased variability in binding conformations during ligand recognition. Together, these results demonstrate that individual TPR repeats are not functionally equivalent and specific regions of the TPR domain play disproportionate roles in maintaining structural integrity and ligand-binding behavior.

### Disease-Associated Variants Disrupt OGT Structure and Alter Predicted Binding Interactions

Disease-associated amino acid substitution mutations within the OGT TPR domain produced substantial structural changes despite affecting only a single amino acid residue. Structural alignment of five XLID-associated variants (L254F, A259T, R284P, A319T, and E339G) revealed differences in the magnitude of structural disruption across mutations (Figure 5a).

**Figure 5:**
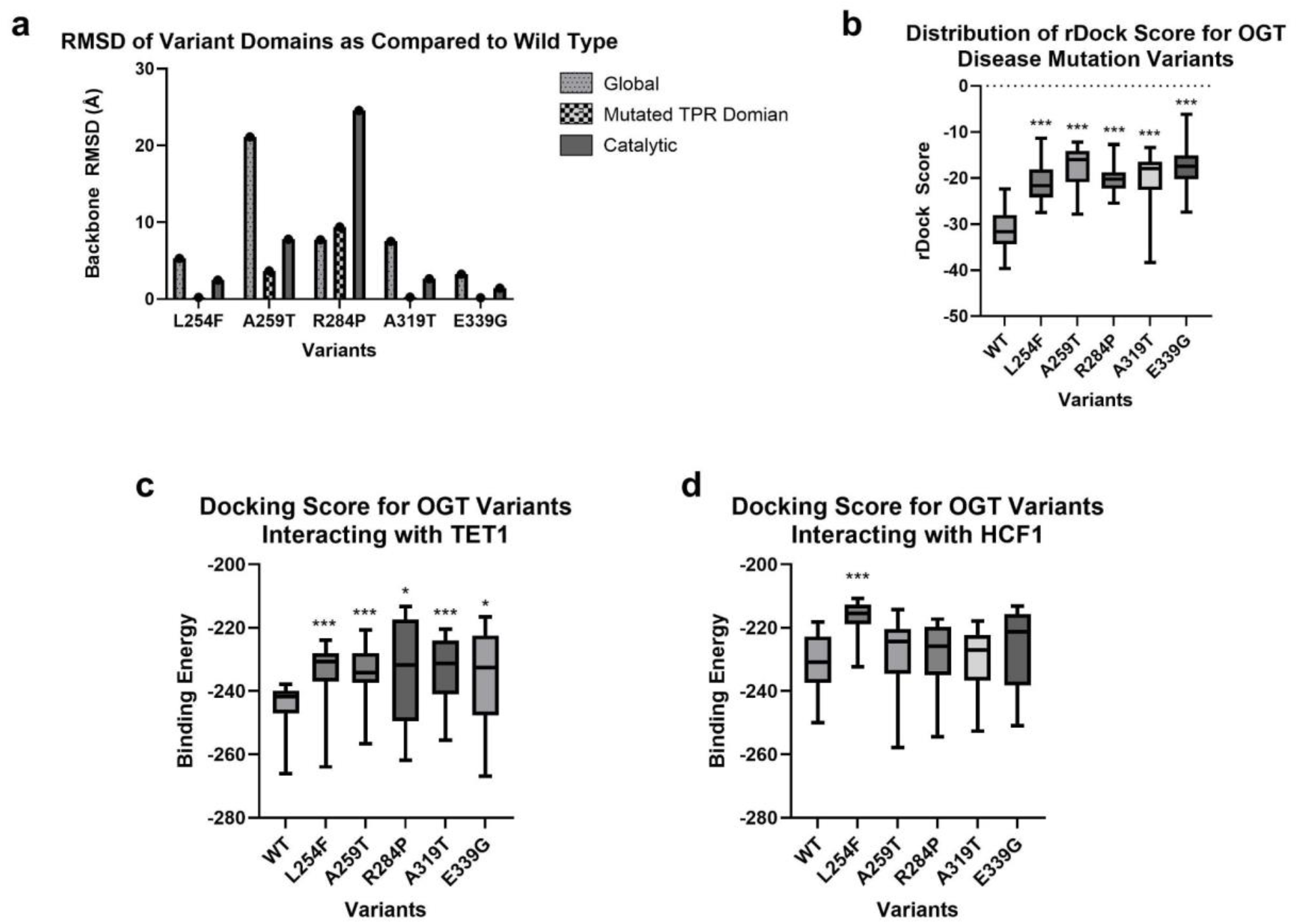
(a) Global, mutated TPR domain, and C-terminal catalytic domain RMSD values for each variant. (b) rDock scores for 20 docking poses (n = 20) of UDP-GlcNAc with wild-type OGT and the selected disease-associated point mutation variants. (c) HDOCK scores for 20 docking poses (n = 20) of TET1. (d) HDOCK scores for 20 docking poses (n = 20) of HCF1. For all docking analyses, more negative scores indicate stronger predicted binding. Statistical significance was determined using pairwise t-tests relative to wild-type (*p < 0.05, ***p < 0.001)

Among the variants, A259T exhibited the largest global structural deviation from wild-type OGT with an RMSD approaching 21 Å, while R284P produced the second-largest deviation at approximately 8 Å. In contrast, L254F, A319T, and E339G resulted in more moderate structural changes.

The effects of these substitution mutations extended beyond the immediate site of the missense mutations. Domain-specific analysis demonstrated that several variants induced substantial conformational changes within the catalytic domain despite mutations being localized to the TPR region (Figure 5a). Notably, R284P produced the largest catalytic-domain deviation, with RMSD values approaching 25 Å. These findings show that local changes within the TPR superhelix can propagate throughout the protein and alter functional regions beyond the TPR domain.

To assess the functional consequences of these structural changes, docking simulations were performed using UDP-GlcNAc. Compared to wild-type OGT, all five variants exhibited significantly less favorable docking scores, indicating reduced predicted ligand affinity (Figure 5b). Although each amino acid substitution weakened predicted binding, the magnitude and variability of these effects differed across variants.

Protein-protein docking analyses further revealed substrate-specific effects of disease-associated amino acid substitutions. All variants had reduced predicted binding affinity for TET1 compared with wild-type OGT (Figure 5c), indicating that disruption of TPR architecture can impair interactions with chromatin-associated binding partners. Interestingly, the R284P variant maintained strong predicted TET1 binding despite substantial structural deviation, suggesting a potential decoupling between structural disruption and binding behavior. In contrast, only L254F had statistically significant differences observed in HCF1 (Figure 5d), although increased variability in docking scores was evident for several other variants.

Collectively, these results show that XLID-associated amino acid substitutions alter OGT structure through catalytic domain conformational effects and differentially impact ligand and protein binding interactions. The observed reduction in TET1 binding but relative preservation of HCF1 interactions suggests that disease-associated mutations may selectively rewire protein-protein interactions rather than uniformly abolishing protein function.

### Engineered Isoforms Rewire Protein-Protein Interactions

To determine whether structural perturbations within the TPR domain selectively influence OGT protein-protein interactions, docking simulations were performed between wild-type OGT and a subset of selected variants (delTPR1, delTPR3, delTPR5, and OGT-8) using the biologically relevant interaction partners TET1 and HCF1. These variants were selected to represent distinct regions of the TPR superhelix while preserving the overall catalytic structure of the protein.

Interactions with TET1 were significantly affected by TPR deletions (Figure 6a). Relative to wild-type, both delTPR1 and OGT-8 exhibited significantly altered docking scores, indicating reduced predicted binding affinity. In contrast, delTPR3 and delTPR5 had binding scores comparable to wild-type OGT, suggesting that these regions contribute less substantially to TET1 recognition. Together, these findings suggest that specific regions of the TPR superhelix are required to maintain productive OGT-TET1 interactions.

**Figure 6:**
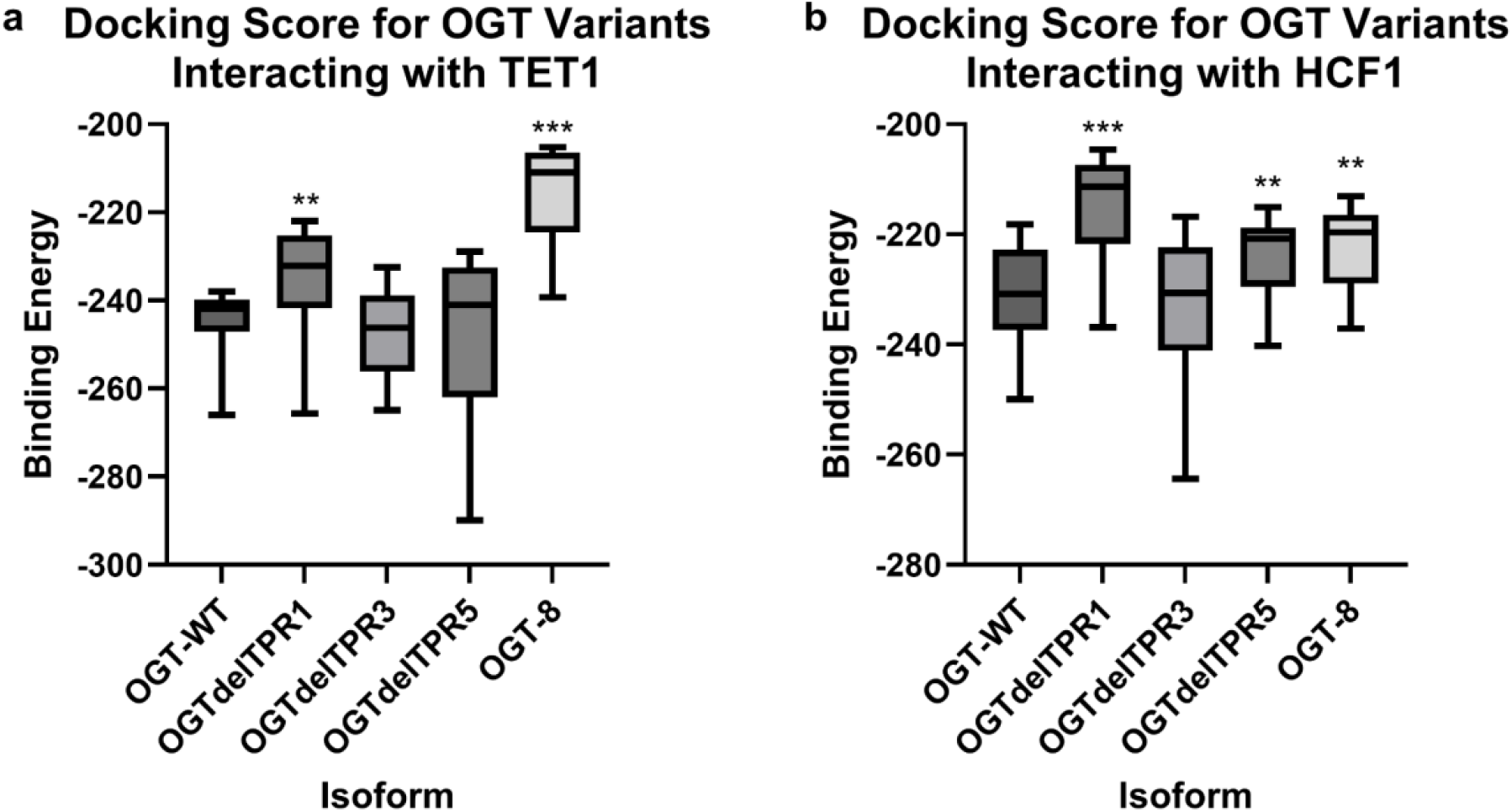
(a) HDOCK scores for the top 20 poses (n = 20) between TET1 and wild-type OGT, TPR deletion variants, or naturally occurring isoforms. (b) HDOCK scores for the top 20 poses (n = 20) between HCF1 and wild-type OGT, TPR deletion variants, or naturally occurring isoforms. For all docking analyses, more negative scores indicate stronger predicted binding. Statistical significance was determined using pairwise t-tests relative to wild-type. *p < 0.05, **p < 0.01, ***p < 0.001.

Distinct interaction patterns were also observed for HCF1 (Figure 6b). Compared with wild-type OGT, delTPR1, delTPR5, and OGT-8 exhibited less negative docking scores, whereas delTPR3 was statistically indistinguishable from wild-type. Unlike the broader score distributions observed for TET1, HCF1 docking produced narrower distributions across all variants, indicating more stable predicted binding conformations. The sensitivity of HCF1 protein interactions to deletion of specific TPR domains suggests that multiple regions of the TPR superhelix contribute to substrate recognition and positioning.

Comparing TET1 and HCF1 docking results revealed that variants that significantly altered binding to one substrate often produced minimal effects on the other, indicating that individual TPR perturbations do not uniformly affect all interaction partners. Taken together, the interaction analysis demonstrates that targeted engineering of TPR domains can modify OGT protein-protein interactions with disruptions to the overall protein structure. This ability highlights the potential of repeat-domain engineering as a strategy for requiring protein interaction networks to alter substrate recognition.

## Discussion

This study supports the idea that the TPR domains of OGT influence both overall protein conformation and interaction specificity with other proteins. Analysis of previously published synthetic isoforms revealed that both excessive truncation (OGT-4) and relative extension (OGT-8) of the TPR array caused the biggest global and catalytic-domain RMSD changes, while intermediate-length variants remained more similar to wild-type^13,14^. However, the structural effects of sequential TPR deletions were not uniform across all isoforms. OGT-6 had similar structural deviation in the TPR and catalytic regions, while OGT-7 showed greater deviation within the TPR region. The limited structural impact on the catalytic domain seen in these isoforms suggests that the structural consequences of TPR modification may not only depend on how many TPR domains are removed, but also on which TPR domains are kept. Certain TPR deletions may better preserve the organization of the catalytic domain despite alterations being made to the TPR superhelix. Overall, this pattern suggests the existence of an optimal TPR domain length required to maintain proper protein conformation, which matches prior work showing that isoforms with fewer than eight TPRs compromise OGT function and cell viability^5^. The high RMSD value observed for the catalytic domain of OGT-4 and OGT-8 further shows that alterations within the TPR region can be associated with structural changes propagating to the C-terminal catalytic domain, even though the catalytic domain amino acid sequence was unchanged. This supports a potential long-range structural relationship between the TPR superhelix and catalytic domain.

Analysis of single TPR deletion variants demonstrated that individual repeats contribute unequally to structural stability. Deletions in early TPRs produced minor increases in global and catalytic-domain RMSD, whereas deletions in mid-to-late TPRs (TPR9-TPR13) resulted in larger structural deviations and higher catalytic-domain RMSD, indicating that these regions play a critical role in maintaining the integrity of the superhelical architecture^15^. In most variants, structural deviations were more pronounced within the TPR region than the catalytic domain, suggesting that structural perturbations are often localized near the deletion site, but not confined to that location. Ligand docking results further demonstrated that central TPR deletions were associated with reduced binding affinity, as indicated by less negative rDock scores, while terminal deletions caused more variable score distributions. These findings suggest that distinct regions of the TPR domain may control substrate accessibility and binding pocket stability, as has been shown for TPR domains of protein phosphatase 5^16^.

Comparisons between wild-type OGT and disease-associated missense variants show that although all the mutations were localized to the TPR domain, they produced disproportionately large effects within the catalytic domain. Perturbations in the TPR region can propagate to distal regions of the protein, influencing catalytic-domain conformation despite no direct sequence modification in that region. Consistent with this observation, no statistically significant differences in HCF1 binding were detected across the point mutations, indicating that structural disruption does not uniformly translate to altered protein-protein interaction strength for this substrate. Further investigation with additional structural metrics should be leveraged to examine this. Together, these findings suggest that while amino acid substitutions in the TPR domain are associated with disease phenotypes, their functional impact in some cases may arise from a destabilized catalytic domain rather than direct disruption of TPR-mediated interactions^3^.

Docking to TET1 and HCF1 connects these structural changes to protein-protein interactions that matter for XLID. Variants such as delTPR1 and OGT-8, which exhibited larger structural deviations, also showed significantly shifted HDOCK scores for one or both partners, whereas deletions such as delTPR3 or delTPR5 showed minimal or substrate-specific effects. These observations are consistent with clinical data indicating that disease-associated variants in OGT cluster in specific TPR regions and can selectively interfere with HCF1 processing or chromatin-associated complexes without completely knocking out OGT activity^3^. Collectively, these results support a model where certain segments of the TPR domain encode preferred binding surfaces for specific proteins, such that structural perturbations can rewire protein-protein interaction networks rather than simply eliminating function.

Several limitations should be considered when interpreting these findings. All analyses were conducted using computational structure prediction and docking (Boltz, AlphaFold3, Speak to a Protein, and HDOCK), which may not capture dynamic conformational changes, post-translational modifications, and cellular factors that influence OGT behavior in real cells. Additionally, RMSD and docking scores provide indirect measures of structural and functional impact. Increased RMSD does not necessarily correspond to a loss of function, and more negative scores do not guarantee stronger binding or correct catalysis. The use of multiple modeling tools may also introduce small tool-dependent biases.

Continuing use of AI tools is important to the further development of predictive modeling of proteins. Our experience with Speak to a Protein demonstrates the potential for protein-focused AI co-scientists for accelerating protein structure analysis and interpretation using natural language. However, many changes are needed before such approaches can be reliably used. In this study, we encountered numerous issues with back-end processing due to the high volume of requests to protein structure inference tools, which led to delays in the analysis. Further, during the process, we noted multiple times when the protein co-scientist tool would claim to have completed a task; however, upon manual inspection, the tasks were not complete. Issues like these need to be addressed before the routine use of these tools.

Despite these limitations, combining modeling of truncated TPR variants of OGT, single-TPR deletions, and disease-associated variants with docking aided by several AI tools gives a systematic framework for identifying structurally and functionally critical regions within the OGT TPR domain. Future work should expand this approach by examining less probable models of each variant to capture alternative binding poses that may be functionally relevant, as well as incorporating molecular dynamics simulations to better understand binding behavior over time. Experimental validation of high-impact variants will also be essential, including biochemical assays to measure changes in glycosylation activity and substrate binding. More broadly, this study demonstrates that the integration of a protein co-scientist with protein structure simulation tools and multimodal AI tools can be used to predict and engineer protein isoforms with altered substrate binding and protein-protein interaction profiles.

## Notes

### Competing Interest Statement

The authors have declared no competing interest.

